# What the flip? What the P-N flip can tell us about proactive suppression

**DOI:** 10.1101/2022.03.04.483004

**Authors:** Joyce Tam, Chloe Callahan-Flintoft, Brad Wyble

**Author notes:** Correspondence concerning this article should be addressed to Joyce Tam, Department of Psychology, Pennsylvania State University, University Park, PA 16801. **Author Note**.

## Abstract

It has been debated whether salient distractors in visual search can be proactively suppressed to completely prevent attentional capture, as the occurrence of proactive suppression implies that the initial shift of attention is not entirely driven by physical salience. While the presence of a Pd component in the EEG (associated with suppression) without a preceding N2pc component (associated with selection) has been used as evidence for proactive suppression, the link between these ERPs and the underlying mechanisms is not always clear. This is exemplified in two recent papers that observed the same waveform pattern, where an early Pd-like component flipped to a N2pc-like component, but provided very different interpretations (Drisdelle & Eimer, *Psychophysiology, 58*(9), 1-12, 2021; Kerzel & Burra, *Journal of Cognitive Neuroscience, 32*(6), 1170-1183, 2020). Using RAGNAROC (Wyble et al., *Psychological Review, 127*(6), 1163-1198, 2020), a computational model of reflexive attention, we successfully simulated this ERP pattern with minimal changes to its existing architecture, providing a parsimonious and mechanistic explanation for this flip in the EEG that is unique from both of the previous interpretations. Our account supports the occurrence of proactive suppression and demonstrates the benefits of incorporating computational modelling into theory building.

---

A major debate in the study of reflexive visual attention is whether or under what conditions attention would be captured by distractor items and such capture can be minimized or perhaps avoided entirely through suppression. Most impressively, it has been demonstrated that some cases of distractor suppression can occur so early that distractor-related attentional capture is completely prevented (see Geng, 2014; Luck et al., 2021). This type of preventive suppression may occur as *proactive suppression*, in which attributes (feature or location) consistently associated with the distractor are inhibited *prior to the onset of the search display* (as suggested by the adjusted version of the signal suppression theory, SST, Gaspelin & Luck, 2019) ^1^.

The stimulus-driven selection account (Theeuwes, 1992, 2010), among others, suggests that the *initial shift* of attention after display or fixation onset is solely driven by bottom-up saliency signals in an automatic manner, such that attention is first attracted to the highest-salience location. Under this theory, attentional capture cannot be prevented, as a salient distractor can be suppressed reactively only after it has captured attention. On the other hand, some theories have argued for an early role of top-down control that would allow for the initial attentional shift to deviate from the saliency signals and thus allow for the occurrence of proactive suppression. For example, the contingent-capture theory (Folk et al., 1992) suggests that salient distractors capture attention only when they are contingent with the task set. Furthermore, the signal detection theory (SST, Sawaki & Luck, 2010; Gaspelin & Luck, 2019) directly suggests that the attraction signals (“attend-to-me signals”) from the salient distractors can be prevented by suppression mechanisms.

In their representative works on SST, Gaspelin et al. (2015, 2017) found evidence for proactive featural suppression when the target and the salient distractor each has a consistent feature (we call it “fixed feature search” in this article). For example, in Experiments 2, 3 and 4 in Gaspelin et al. (2015), participants searched for a pre-defined shape target among heterogeneous shapes. In half of the trials, one of the distractors had a color different from the rest of the items. This distractor color was held constant across the experiment, and thus the participants were able to learn the color association. No attentional capture effect was found, in that the presence of a color singleton did not slow reaction times to the target. Furthermore, to measure the spatial distribution of attention, letter probes appeared briefly on the search items in about 30% of the trials, and participants were to report as many letters as they could. Critically, they observed the *probe suppression effect*, i.e., the letter probe at the singleton location was reported *less accurately* than probes at nonsingleton distractor locations, such that the singleton’s location was the least attended to among all items (see also Chang & Egeth, 2019, 2021). Importantly, the probe suppression effect was observed even for very briefly presented probes (100ms) with zero display-to-probe delay (e.g., Experiment 4 in Gaspelin et al., 2015), making a rapid disengagement account (Theeuwes et al., 2000; Theeuwes, 2010) improbable. This effect was found even with a highly salient distractor (a singleton color in a display of 20 items) with its stand-out saliency confirmed by an independently computed estimation (Stilwell & Gaspelin, 2021; Stilwell et al., 2022).

In addition, with electroencephalogram (EEG) data, the relative polarity between parietal-occipital scalp sites contralateral and ipsilateral to a visual item of interest (typical sites are PO7 and PO8) has been found to correlate with attention. Typically observed at around 200ms from stimulus onset, a contralateral negativity, or the N2pc, correlates with attentional selection (Luck & Hillyard, 1994); whereas a contralateral positivity, or the Pd, correlates with attentional suppression (Hickey et al., 2009). These two components have been adopted as tools to measure these attentional mechanisms with a particular emphasis on their temporal sequencing. For example, a Pd component towards the singleton distractor has been observed without a preceding N2pc, suggesting that the suppression process can occur proactively (Gaspar & McDonald, 2014; Gaspelin & Luck, 2018a, Jannati et al., 2013; Sawaki & Luck, 2010; Stilwell et al., 2022).

Although the N2pc and the Pd have played pivotal roles in the understanding of attentional suppression, we must keep in mind that there is no way to distinguish between attention of the item(s) on one side and suppression of the item(s) on the other side solely based on the EEG signals. This issue is exemplified in a recently observed ERP sequence (Drisdelle & Eimer, 2021; Kerzel & Burra, 2020). In detail, Kerzel & Burra (2020) suggested that at least some of the evidence for proactive featural suppression, especially those coming from small displays (but see Stilwell & Gaspelin, 2021; Stilwell et al., 2022), could be explained by a form of serial search when the items were heterogeneous (but see Gaspar & McDonald, 2014; Jannati et al., 2013). To investigate, Kerzel & Burra (2020) replicated the general experimental design used in Gaspelin et al. (2015) and analyzed EEG signals from lateral parietal-occipital sites. They observed an early Pd-like contralateral positivity (∼185-235ms, relative to the singleton distractor location), replicating Gaspelin & Luck (2018a). However, Kerzel & Burra (2020) also observed an N2pc-like contralateral negativity trailing the initial positivity (∼265-315ms), forming what we term here the P-N flip^2^. As it would not make sense for the attentional system to re-attend to a distractor that had already been suppressed, Kerzel & Burra (2020) proposed that the ERPs were in fact two successive N2pc components – the first one reflecting attention towards the lateral nonsingleton distractor, the second one attention towards the salient distractor. This might have occurred due to the use of a specific serial scanning strategy afforded by those specific task settings (small set size, heterogeneous displays), with three additional assumptions: (1) participants tend to scan lateral items prior to vertical items, (2) when the salient distractor is one of the lateral items, they tend to scan the opposite-side distractor first because the consistent salient distractor color forms a “template for rejection” (see Arita et al., 2012) – leading to an initial attentional enhancement of the lateral nonsingleton distractor, and (3) they are unable to skip scanning the salient distractor after that, despite this template for rejection – leading to the subsequent attentional enhancement of the salient distractor. Note that Kerzel & Burra (2020) also emphasized that these interpretations were primarily speculative. However, if their interpretation is correct, it would mean that the presence of a consistent distractor feature may not prevent capture by singletons so much as delay it under some circumstances.

According to this serial search hypothesis, the P-N flip should disappear when there are no incentives for the participants to attend to the lateral positions. To test this idea, Drisdelle & Eimer (2021) included a condition in which the target never appeared laterally (the “focused midline” condition). Although participants were able to utilize this spatial regularity to their advantage, as they were substantially faster to respond to the target compared to the “unfocused” condition (where the target could appear at any of the four locations), the P-N flip was reliably observed in both conditions – meaning that a lateral-first serial search strategy cannot completely explain the P-N flip. Drisdelle & Eimer (2021) in turn proposed that the ERPs were two successive Pd components – the first one reflecting suppression of the singleton distractor, and the second one reflecting suppression of the lateral nonsingleton distractor. They further suggested that the suppression of the singleton distractor occurred earlier because of its superior salience. This is in line with the proposal of *active suppression* based on relative saliency signals (Sawaki & Luck, 2010; see also footnote 1). However, this proposal is subject to challenges from studies that fail to observe successful preventive suppression of singleton distractors without a predictable feature value (e.g., Gaspelin & Luck, 2018b). This point will be further discussed in a later part of this article.

Motivated by this puzzle, we tried to simulate the P-N flip with RAGNAROC (Wyble et al., 2020). By modelling the dynamics of interacting neural populations, RAGNAROC provides a mechanistic description of the selection and suppression processes in covert reflexive attention, as well as how these underlying sources are linked to the associated ERPs (i.e., N2pc and Pd). Note that RAGNAROC was not designed with the explicit purpose to simulate the flip. It would therefore provide a parsimonious account if the simulation can be achieved with little to no alterations to the model.

With the model, we implemented a proactive form of distractor suppression that would be effective when the feature values of a distractor were consistent across trials. This was done by reducing the connection weights of features that are consistently and specifically associated with singleton distractors. With this change, RAGNAROC successfully simulates the flip using its existing circuitry (Figure 1). This provides a perspective on the underlying cause of the P-N flip that is unique from Kerzel & Burra (2020) and Drisdelle & Eimer (2021). We further explored the effect of target presence, set size, and the inclusion of probe trials on the simulation results in a fixed feature search, referencing relevant empirical evidence (Barras & Kerzel, 2016; Drisdelle & Eimer, 2021; Gaspelin et al., 2015; Gaspelin & Luck, 2018a; Stilwell et al., 2022). All in all, our account supports the possibility of the occurrence of proactive featural suppression and shows *how* this would lead to the pattern of waveforms that have been observed. It also acts as an example to demonstrate some of the advantages in incorporating computational modelling into theory building. In the next section, we will introduce the model’s architecture and illustrate the way in which these simulations were implemented.

**Figure 1.**
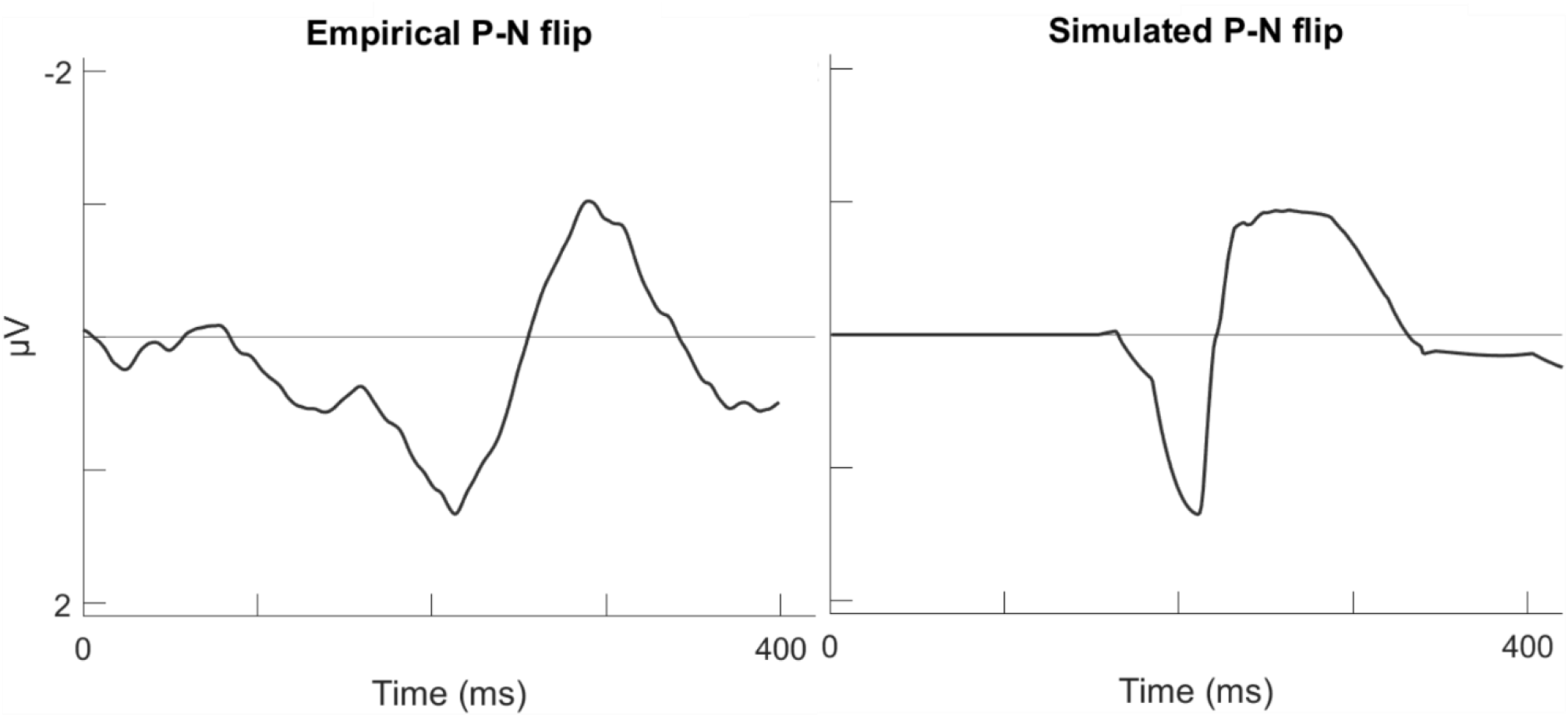
RAGNAROC’s simulation of the P-N flip in fixed feature search. Both the empirical and simulated EEGs were generated from a target midline, singleton distractor lateral configuration with a set size of 4. Plotted are the difference waveforms between contralateral and ipsilateral EEG signals at occipital-parietal sites, pivoting on the singleton location. The empirical P-N flip in Kerzel & Burra (2020)’s experiment 2 (left) is simulated by RAGNAROC (right).

## Model Mechanisms Underlying the P-N Flip

While the explanations from Kerzel & Burra as well as Drisdelle & Eimer focus on an iteration of attentional processes between the lateralized nonsingleton distractor and the salient distractor, our work with RAGNAROC proposes an important alternative in which the consistent singleton distractor has essentially no effect on attention because the visual attention system has learned to reduce its top-down relevance weighting to nearly zero. As a result, both the P and N components are instead caused by the nonsingleton distractor on the other side.

In order to explain this, we will first provide a summary of the model, although readers are suggested to refer to Wyble et al. (2020) for a thorough introduction. Figure 2 shows RAGNAROC’s macro-architecture as sheets of simulate neurons that correspond to stages of visual processing. Visual information presented to RAGNAROC passes through three layers of retinotopically arranged neurons in this order: early vision (EV), late vision (LV) and attention map (AM). The model assumes that physical saliences of stimuli are computed in the EV and affect the strength of signals projected to the feature-specific maps in LV. The next step from LV to AM incorporates top-down relevance weightings corresponding to the current goals of the observer. Taking into account the interaction of the physical saliences and relevance weightings, the AM computes where attention will be deployed and where it will be suppressed. Importantly, the dynamics of the AM activations are translated into EEG recorded at posterior-occipital sites, with higher AM activations resulting in more negative EEG signals on the contralateral side. The mechanisms of RAGANROC are supported by its ability to simulate behavioral and electrophysiological data from an array of empirical benchmarks (see Wyble et al., 2020).

**Figure 2.**
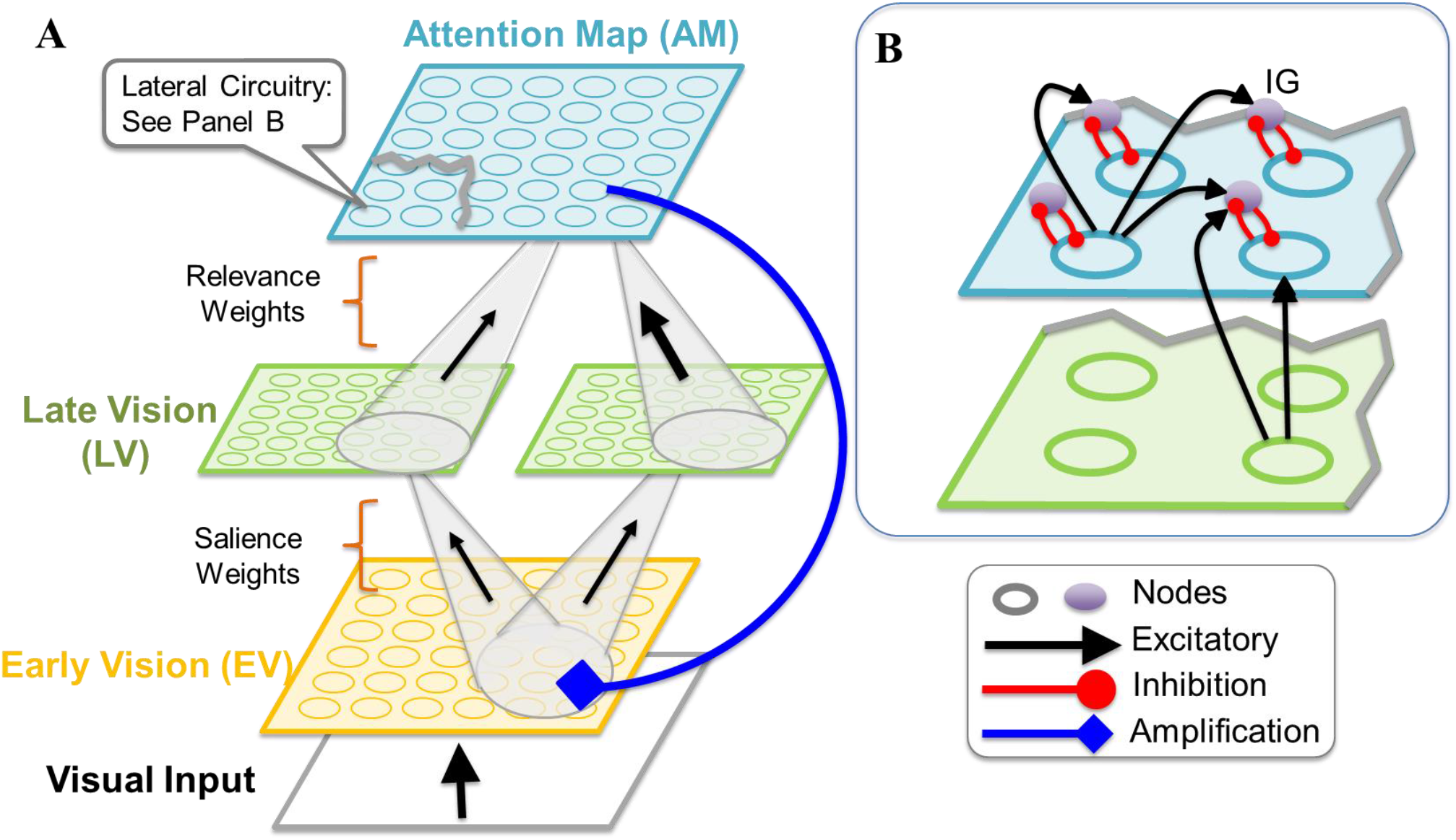
The macro-architecture of RAGNAROC adapted from Figure 4 in Wyble et al. (2020). Visual input passes through EV, LV, and finally the AM. The saliency signals are represented in connections from EV to LV. Then, top-down relevance weightings can vary to modulate signals from the feature-specific LV maps, indicated by the different thicknesses of the arrows from LV to AM. The down-weighting of the distractor-associated feature is the basis to implement proactive featural suppression. Panel B shows how *reactive* suppression is implemented only for items with sufficient AM activation. Therefore, a proactively suppressed salient distractor receives little-to-no reactive suppression while the nonsingleton distractors that are not proactively suppressed receive stronger reactive suppression. These mechanisms combined allow RAGNAROC to successfully simulate the P-N flip.

The important aspect of RAGNAROC as regards the P-N flip is the competitive inhibitory circuits that implement a reactive form of inhibition that is *proportional to stimulus strength*. As signals first reach the AM, a brief competition across the entire AM occurs to determine which regions are to be attended to. When the competition is resolved, the “losers” (i.e., areas with weaker activation) will be suppressed in proportion to the input they receive from the LV, i.e., *a more activated item will be more deeply inhibited if it loses the competition for attention*. This mechanism is informed by behavioral tests showing that locations containing distractors are more inhibited than blank regions (Cepeda et al., 1998). In RAGNAROC, this same inhibitory circuit is also thought to be the source of selective inhibition of salient distractors without a consistent feature value (Burra & Kerzel, 2013; Kiss et al., 2012; McDonald et al., 2013). We also note here that this reactive inhibitory circuit is spatially graded: Locations closer to the winning stimulus will be more strongly suppressed. This point will be revisited when the effect of set size on the EEG is considered in a later section.

Now consider a case of fixed feature search, where a color singleton distractor is always red, and the target never appears in red. In RAGNAROC, proactive suppression of this consistent distractor feature can be implemented by reducing the feature-specific relevance weighting from the LV to the AM. In turn, the ability of a location containing that feature to recruit attention can be greatly reduced, even if it contains a highly salient item. Thus, the input to the AM driven by a consistent singleton distractor can be *weaker* than input driven by nonsingleton distractors that are not proactively suppressed. The mechanisms in the model dictates that a location with a weaker AM input will translate into a more positive EEG signal at the contralateral electrode, and thus, in a contralateral subtraction, results in a positivity. This simulates the first component in the P-N flip.

The dynamics of stimuli activations from the EV to the AM in the case of proactive suppression is further illustrated in Figure 3. Importantly, this implementation of proactive suppression cannot suppress an item below zero on the AM. Instead, the AM activation will be at approximately zero, and the AM dynamics will develop as if the distractor was absent. Thus, proactive featural suppression can be better described as a strong “ignoring” of the item through feature downweighing. However, the salient distractor is “suppressed” in the sense that this feature value’s weight is selectively inhibited.

**Figure 3.**
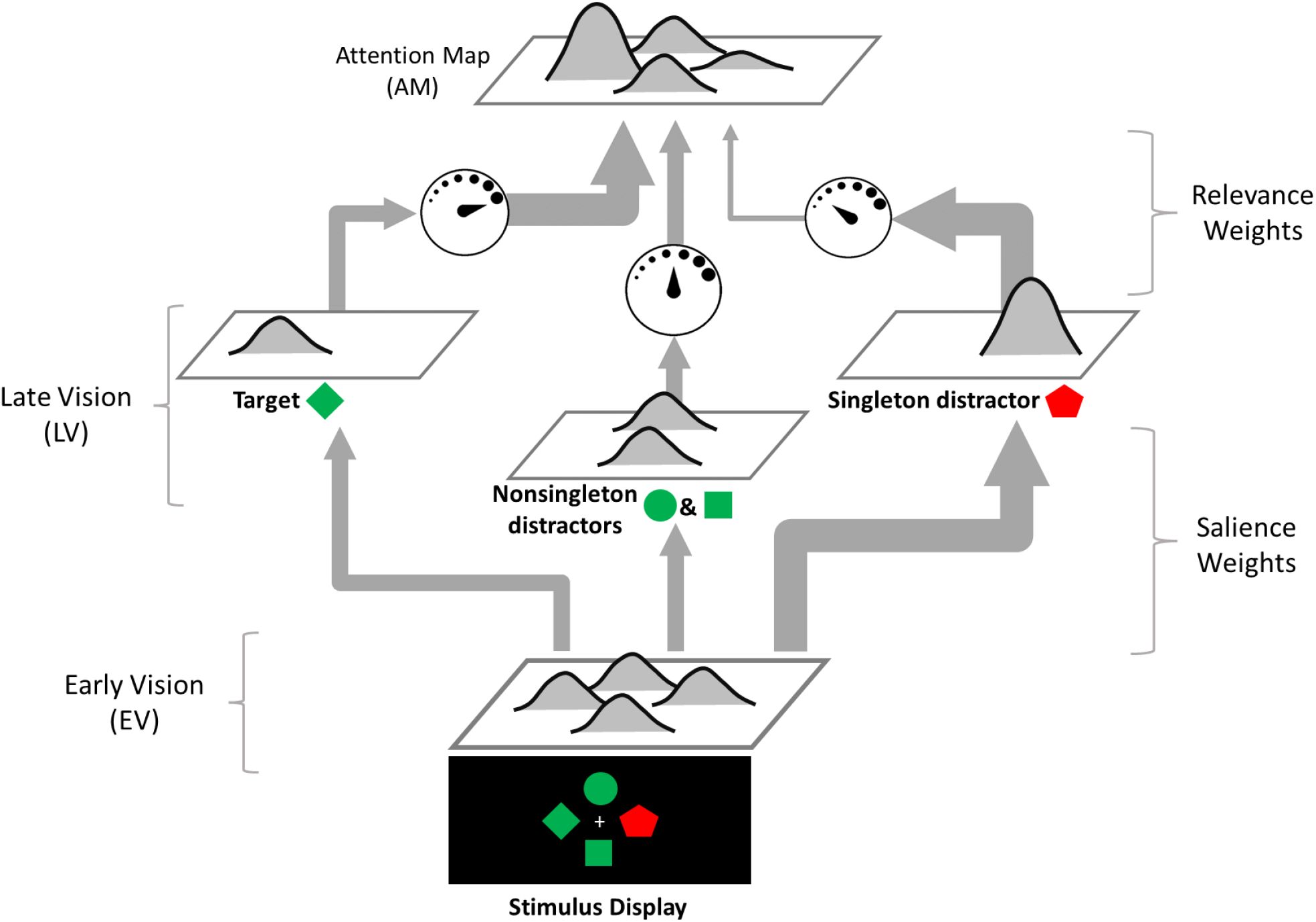
The implementation of proactive suppression in RAGNAROC. In this example, participants always searched for a green diamond and one of the distractors always appeared in red, allowing it to be proactively suppressed. The singleton distractor is the most salient, thus producing the strongest LV input as shown by the thickness of the arrows from EV to LV. However, the neural activities are then modulated by task relevance weights before entering the AM. The relevance weights are indicated by the dials in the top half of the figure. The diamond shape is associated with the target and is therefore “dialed up”. The color red, on the other hand, is associated with the singleton distractor and is therefore “dialed down”, indicating proactive suppression of this input. As a result, in the AM, the target became the most activated, followed by the nonsingleton distractors whose corresponding signals were unmodulated. The singleton distractor location ends up with the weakest AM signals. When the same target is placed at the horizontal midline, the mechanisms illustrated in this figure contributes to the EEG dynamics in the period where the P component in the flip can be observed (see blue boxes in Figure 4).

Given that the proactively inhibited singleton distractor produces an extremely small AM signal, how does it then appear to elicit a subsequent N component that can be interpreted as attention towards itself (Kerzel & Burra, 2020)? To this point, RAGNAROC provides a subtle and interesting alternative to understand the P-N flip using the characteristic of *proportional inhibition* in its reactive suppression mechanism (visualized in Figure 4). As the AM input at the singleton distractor location is weaker than that at the nonsingleton distractor location, reactive inhibition from the target is weaker for the singleton location. Thus, as reactive inhibition progresses, the nonsingleton distractor locations get suppressed more and eventually to the point where they are less activated than the singleton distractor location. This results in the singleton location AM activities being *relatively stronger*. The mechanisms in the model dictates that a location with a stronger AM activity will translate into a more negative EEG waveform at the contralateral electrode, and thus, we see a relative negativity after contralateral subtraction. This simulates the second component of the flip (see the green boxes in Figure 4).

**Figure 4.**
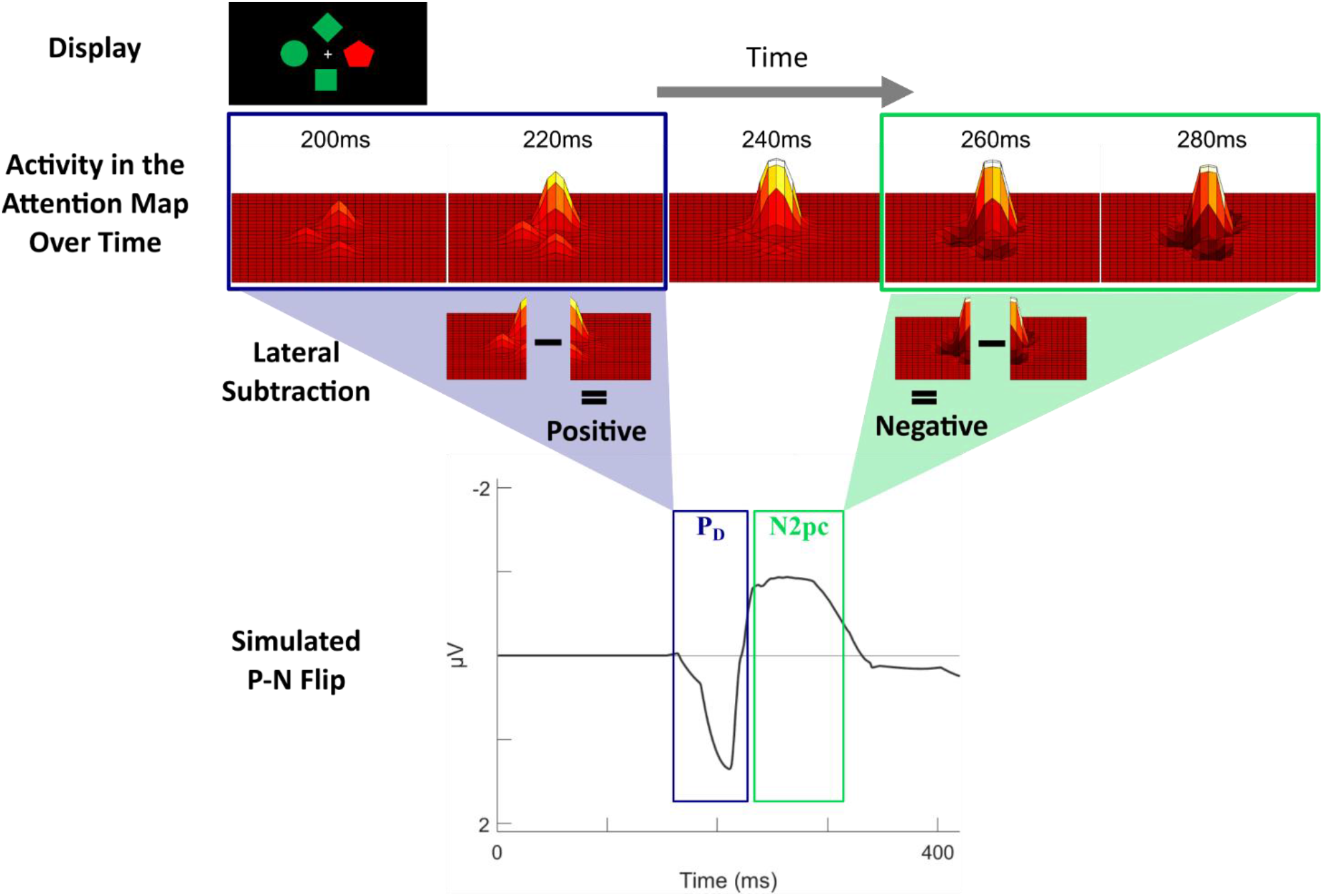
Attentional dynamics that underlie RAGNAROC’s simulation of the P-N flip. In this example, participants always searched for a green diamond and one of the distractors always appeared in red, allowing it to be proactively suppressed. **Top:** Relevant dynamics in the Attention Map (AM) at the critical timepoints for the flip (200 – 280ms after array onset). A positive deflection on the map indicates enhancement of attention for items at the deflection location. The blue box indicates the P component: The lateral nonsingleton distractor (NS) had a weak but unsuppressed activation, whereas the singleton (S) had a near-zero activation due to feature downweighing. Technically, the weaker activation at the singleton location translates to a relative positivity at its contralateral electrode, resulting in a positivity after contralateral subtraction. As the polarity of this resultant component will always be the same as the result of directly subtracting the AM activities of the singleton side from that of the nonsingleton side, the process of contralateral subtraction is visualized as subtraction between AM activities from the two sides for ease of visualization. The green box indicates the N component: Unselected items are suppressed proportional to their activation strengths. As a result, the more-activated NSs got suppressed more than the less-activated S, resulting in a contralateral negativity. **Bottom:** A simulated ERP is generated from the simulated AM activity by just averaging the total synaptic activity for each side of the AM and subtracting one side from the other.

Unlike proactive suppression, this reactive inhibition occurring at the AM will suppress the level of attention at those locations below baseline. This allows for a critical empirical test between proactive and reactive suppression that we will illustrate below.

### Predicted Empirical Distinction of Proactive and Reactive Suppression

In RAGNAROC, proactive and reactive suppression have distinct mechanisms with different effects on behavior and EEG. The most critical difference is how attention is allocated within the AM. Proactive suppression prevents attention towards the distractor by effectively ignoring the distractor. Therefore, it does not suppress the corresponding AM signal below zero. On the other hand, reactive suppression can produce inhibitory wells (Figure 4A, 260-280ms). Thus, *we predict that a proactively suppressed location will never be activated less than an empty location whereas a reactively suppressed location will possibly be*.

The test could be as follows: A salient color singleton with a predictable color is sometimes presented in a visual search task. When the target is presented on the midline and the salient distractor is lateral, we predict an EEG pattern resembling the P-N flip if a nonsingleton distractor occupies the location contralateral to the salient distractor. In contrast, no observable ERPs will be produced if the contralateral side is empty. Again, this is because when the salient singleton has a consistent feature, both the P and N components of the P-N flip are caused by the nonsingleton distractor. It should be noted here that a few studies have inspected the EEG waveforms with respect to a consistent-color distractor positioned contralateral to an empty space, but a Pd was observed (Hickey et al., 2009; Hilimire et al., 2012). However, this Pd occurred slightly later (onset at ∼220ms) than the P component in the P-N flip (∼185ms in Kerzel & Burra, 2020; ∼110ms in Drisdelle & Eimer, 2021) and was sometimes preceded by a N2pc; a distractor-present RT cost was also observed in some cases (Hilimire et al., 2012). These observations suggest that participants may not have employed proactive suppression, despite it having a consistent color. One speculation is that proactive suppression is only employed when a search task is sufficiently difficult, as proactive suppression is absent in studies that used a highly distinctive target-distractor pair (a shape vs a line), required a simple discrimination task based on the target’s outline, and employed a minimalistic search display with only 1 or 2 items. However, these factors influencing task difficulty are not taken into account by RAGNAROC. Further research (in both empirical and modelling terms) would be required to determine how task requirements influence suppression processes.

The predicted waveform is different for a salient distractor that is reactively suppressed, e.g., when it does not have a consistent feature value. Given the same target-midline, distractor-lateral configuration, an N-P flip is predicted, no matter whether the contralateral side consists of a nonsingleton distractor or an empty space. However, the amplitudes of both components would be smaller when the nonsingleton distractor is present, compared to when it is absent, as that item has an initial non-zero activation and will then be reactively inhibited, discounting the activation difference between itself and the salient distractor in both the initial activation and the reactive suppression stages.

### Extensions of the P-N flip Simulation

#### Effect of Target Presence

Whether the search target is present critically affects the competition for attention in the AM. In the example we used to illustrate the P-N flip, the target is assumed to be the “winner” of this competition, and it exerts proportional inhibition towards the nonsingleton distractors (stronger) and the singleton distractor (weaker). However, if the target is absent, the nonsingleton distractors may win the competition, or, no winner is determined, and the AM activations gradually return to baseline. In either case, we will not expect the “flip” because AM signals will not be stronger on the singleton side. Thus, we suggest that target presence selectively influences the N component (see Figure 5A). This is consistent with the data in Drisdelle & Eimer (2021).

**Figure 5.**
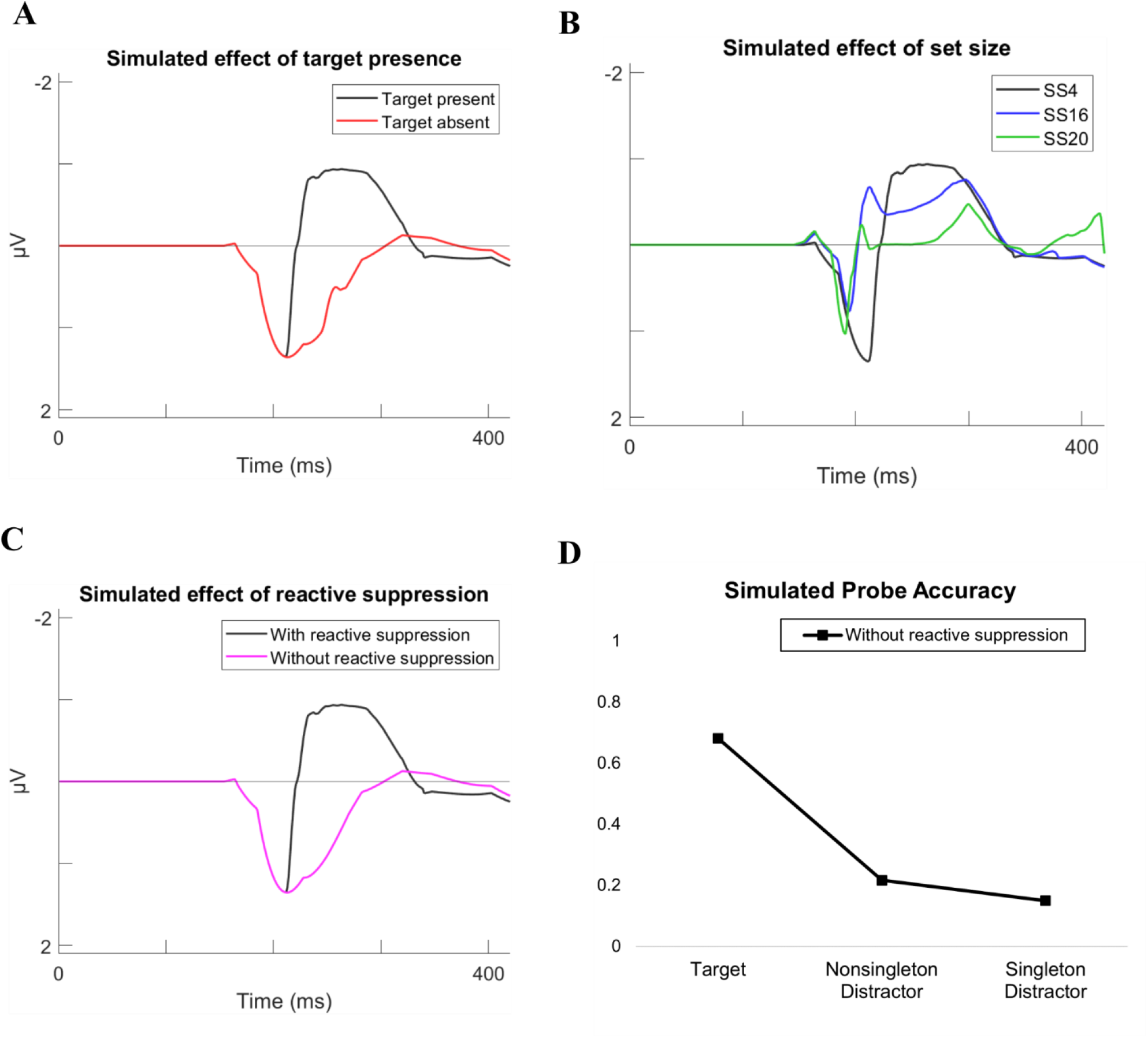
Extending the P-N flip simulation onto other related effects. **(A)** Absence of search targets selectively reduces the N component in the simulation, consistent with Drisdelle & Eimer (2021). **(B)** Increasing the set size changes a number of parameters, including target salience, singleton salience, and inter-item distance. The simulated pattern suggests that the P-N flip is preserved at set size 16, whereas the N component is selectively reduced at set size 20. This mimics the pattern observed in Stilwell et al. (2022). **(C – D)** Simulations performed with reactive suppression disabled in RAGNAROC in order to implement the decrease of spatial selectivity when probe trials are included. **(C)** The N component is selectively reduced, consistent with the visual inspection of Experiment 1 in Gaspelin & Luck (2018a). **(D)** The probe suppression effect observed in Experiment 2 in Gaspelin et al. (2015) can still be successfully simulated when reactive suppression is disabled.

#### Effect of Set Size

The effect of set size on the P-N flip can be complicated and counterintuitive because multiple parameters covary as set size changes. For example, as set size increases, the singleton is thought to be further “singled out”, resulting in a higher salience (Stilwell & Gaspelin, 2021; Stilwell et al., 2022; Wang & Theeuwes, 2020). In addition, increasing the set size often comes with a cost to the display heterogeneity, as shapes are often reused among the distractors (Barras & Kerzel, 2016; Stilwell & Gaspelin, 2021; Stilwell et al., 2022; Wang & Theeuwes, 2020). This makes the target shape more unique and increases the salience of the target. Furthermore, having more items on the screen often means that they are closer together. As noted earlier, reactive suppression in RAGNAROC depends on the graded lateral connections among the AM neurons: Locations closer to the winner will be more strongly suppressed. Therefore, the crowdedness of the stimuli also plays a role in the simulated EEG.

Implementing these parameter changes, we in fact successfully simulated the EEG at set size 16 and 20 reported in Stilwell et al. (2022). At set size 16, the flip is observed, consistent with their Experiment 1. At set size 20, the P component is preserved while the N component is disproportionately reduced or even eliminated, consistent with their Experiment 2 (see Figure 5B)^3^.

#### Effect of Including Probe Trials

In terms of letter probe reports (e.g., Gaspelin et al., 2015), the timing of the P-N flip predicts that only probes presented rapidly after the search array would be suppressed, in contrast to probes presented at a longer latency. However, probe suppression was still observed for probes appearing with a 200ms delay to the onset of the search items (Gaspelin et al., 2015; Gaspelin & Luck, 2018a)^4^.

To account for this inconsistency, it may be important to note that the presence of the letter-probe task may itself affect the mechanisms of attentional selection. In fact, it has been suggested that the inclusion of probe trials influences the way attention is distributed by encouraging the spreading of attention to more item locations, which will in turn influence the EEG (Drisdelle & Eimer, 2021; Kerzel & Burra, 2020). When discussing the findings from Gaspelin & Luck (2018a), Kerzel & Burra (2020) also observed that the inclusion of probe trials in Gaspelin & Luck’s Experiment 1 seems to have selectively eliminated the N component.

In RAGNAROC, a plausible way to simulate the reduction of spatial selectivity of attention is to reduce the connective strengths in the lateral circuits on the AM that mediate reactive suppression. With this assumption, we repeated the simulation of the P-N flip using a version of RAGNAROC where the reactive suppression mechanism is completely disabled. This modification resulted in the selective elimination of the N component (Figure 5C), which is consistent with Kerzel & Burra’s observation. Moreover, this version of the model can still simulate the behavioral probe suppression effect (Figure 5D). Still, although preliminary evidence for the selective influence of probe trials on the N component exists, further studies would be needed to directly assess the influence of the probe trial paradigm or other manipulations of the spread of attention on the EEG in this type of search task.

### Relationships with Existing Observations and Theories

#### One-to-many Mapping from ERPs to Neural Sources

Our account emphasizes that proactive and reactive suppression have distinct mechanisms, which is consistent with the viewpoint of Geng (2014). However, despite having distinct mechanisms and underlying neural sources, RAGNAROC predicts both forms of suppression to result in a positivity for the suppressed item, though at different time points. For a proactively suppressed item, the positivity is caused by the AM bump elicited by the contralateral nonsingleton item, while for a reactively suppressed item the positivity is caused by a stronger suppression of the salient item from the selected item. These simulations are inconsistent with theories that propose a common mechanism for two suppression processes based on them both yielding a Pd (Sawaki et al., 2012), or a seemingly definitive mapping between attention and N2pc, or suppression and Pd (Chelazzi et al., 2019; Gaspelin & Luck, 2018c, Noonan et al., 2018). We suggest instead that the polarity of an ERP alone is not sufficient to drive a strong inference about the underlying causes, especially when the latency differs.

#### Salience May be Unnecessary for Proactive Featural Suppression

Our formulation of proactive featural suppression is essentially an implementation of the down-weighting of a specific feature value. This is largely consistent with the position of the adjusted SST (Gaspelin & Luck, 2019; Gaspelin & Luck in Luck et al., 2021). However, the role of the physical salience of the to-be-suppressed item is often maintained in the adjusted SST, such that an implicit assumption seems to be that the proactively suppressed item must have been a salient distractor to begin with. However, in RAGNAROC, any distractor items can be proactively suppressed if they possess a predictable feature that does not overlap with the target feature. A recent study by Lien et al. (2021) found empirical evidence that distractor saliency is not essential for feature-based suppression to occur. Nonsingleton distractors were introduced in their “triplet” condition, where three items (including the target) consistently shared a color and another three items consistently shared another (distractor) color. As the number of items in the target color and the distractor color were the same, the distractor items were not salient. However, these nonsingleton distractors imposed no cost to search RT and demonstrated a probe suppression effect, indicating that their activations were below that of the distractors sharing the target color.

While we agree with Lien et al. (2021) that saliency is not essential for suppression, in RAGNAROC, singleton distractors do elicit saliency signals (or “attend-to-me signals”) that can lead to attentional capture if not quickly suppressed (Hickey et al., 2006; Theeuwes, 1992, 2004; Theeuwes & Burger, 1998). Also, the idea that saliency is not necessary for suppression does not mean that saliency plays no role in suppression. For example, we speculated earlier in this paper that proactive suppression may only be implemented when a task is sufficiently difficult, making it beneficial to suppress a distractor. In addition, some level of distinguishability is likely required for this distractor-associated feature to be recognized, learnt, and suppressed.

Still, this viewpoint that dissociates proactive featural suppression from physical salience has theoretical implications on the understanding of *active* and *proactive suppression*. To elaborate, Drisdelle & Eimer (2021) interpreted the P-N flip as two acts of suppression. They suggested that the first suppression (towards the salient distractor) occurred earlier due to this distractor’s “attend-to-me signal” driven by its higher relative saliency (see also Sawaki & Luck, 2010). This salience-based suppression mechanism can occur only after the relative physical saliences are computed and thus cannot occur “proactively” (before the stimuli appear). However, as saliency signals are available before the signals are sent to the AM, a successful salience-based suppression may also *prevent* any attention towards the suppressed item, such that it should not be considered an instance of “reactive” mechanisms. Consistent with the original SST (Sawaki & Luck, 2010), we think that this salience-based suppression mechanism should be termed “active” and be distinguished from both proactive and reactive suppression. Whether active saliency suppression actually takes place is a topic for future research.

#### Proactive Featural Suppression as a Form of Template for Rejection

This proactive suppression mechanism can be viewed as a subclass of the idea of templates for rejection (Arita et al., 2012; Carlisle & Nitka, 2019). Arita et al. (2012) reported that precueing a nonsingleton task-irrelevant color sped up visual search, even when the precued color varied randomly from trial to trial. With a different search task, Cunningham & Egeth (2016) also observed a distractor color precueing benefit for search items with heterogenous colors (i.e., the cued color was not salient), but only in the latter half of the experiment when the precued color was consistent throughout. Conceivably, under different circumstances, a template for rejection may be updated based on trial-to-trial cueing (Arita et al., 2012); require repeated use of the same cue (Cunningham & Egeth, 2016); or require repeated exposure to a consistent association between a feature value and the distractor (which is the condition simulated in the current paper).

## Conclusion

In this paper, we described a new account for the P-N flip in fixed feature search (Drisdelle & Eimer, 2021; Kerzel & Burra, 2020) through a combination of proactive featural suppression and reactive suppression implemented with RAGNAROC. We suggest that as the consistent feature of the singleton distractor is proactively suppressed, the singleton distractor is essentially ignored by the attentional system. Thus, the P-N flip is driven by attentional dynamics corresponding to the contralateral nonsingleton distractor, which is first activated, then reactively suppressed. We proposed empirical tests to evaluate this prediction and explored whether the P-N flip simulation is able to account for the empirical effects of target presence, set size, and the inclusion of probe trials on the EEG pattern. In general, we observe a good qualitative compatibility between the data and simulations across these different manipulations.

It should be noted that this paper does not contain a complete mechanistic taxonomy of all possible suppression mechanisms. For example, the mechanism of proactive spatial suppression (e.g., Wang et al., 2019) was not described. Providing a complete mechanistic taxonomy is not our intention. Instead, we use the unintuitive P-N flip to demonstrate how computational modelling can aid the interpretation of empirical data. All empirical data suffers from the reverse inference problem. In the case of N2pc and Pd, which are measured through a lateralized subtraction, the polarity of N and P can be equally caused by a positivity on one side or a negative on the other. A straightforward interpretation of contralateral negativity as attention and contralateral positivity as suppression is thus challenging. The modelling approach allows for formalized theorization of the suppression mechanisms in play and simulations of the ways they manifest, which can be helpful in resolving apparent conflicts between observations and theories, theory choosing, and generating new empirical predictions.

## Appendix

### Parameters

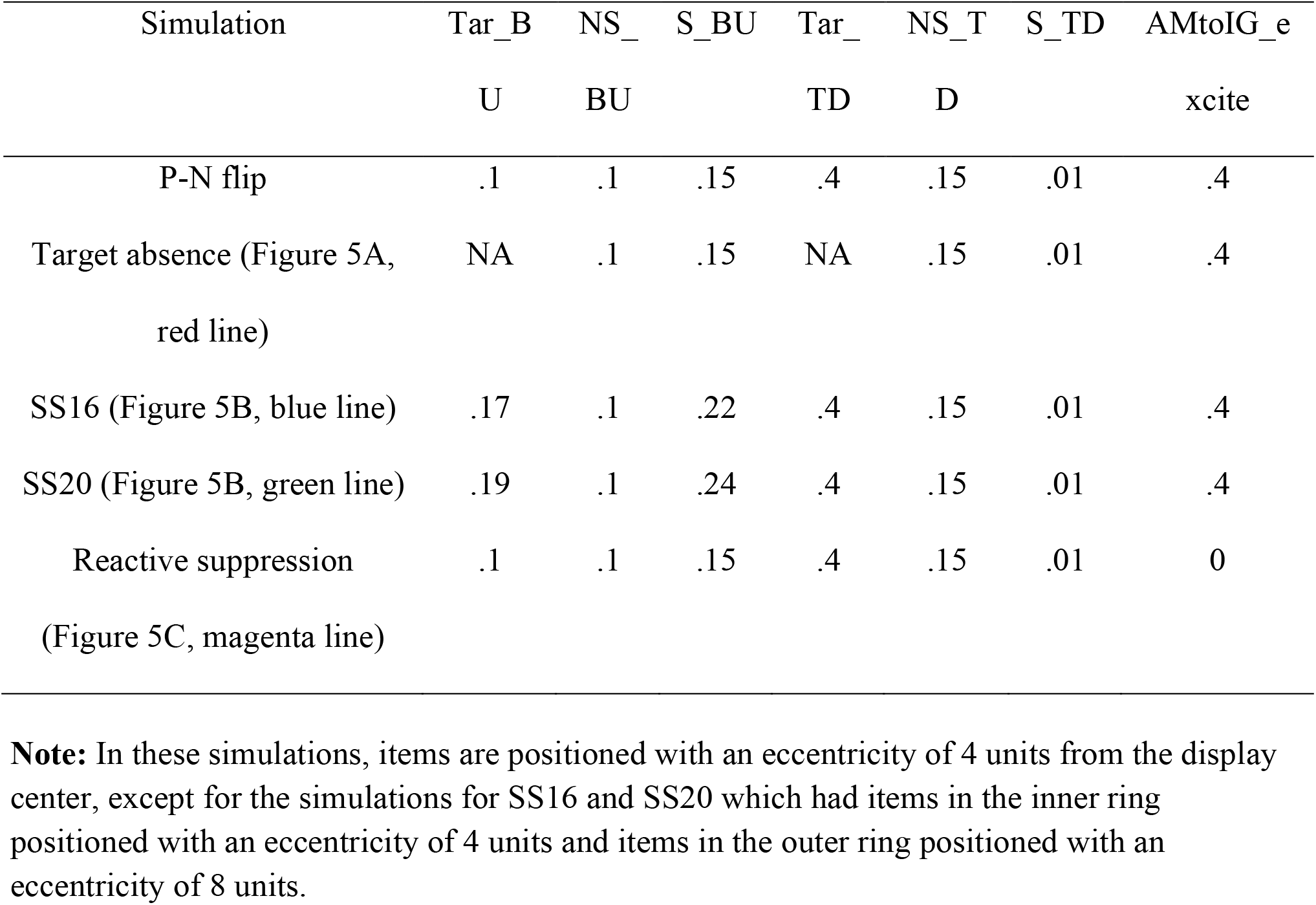

## Footnotes

1 Preventive suppression may also manifest as active suppression, where suppression is said to operate on early saliency signals but only after display onset (Sawaki & Luck, 2010). We note here that this type of salience-based active suppression is sometimes discussed as a mechanism of “proactive” suppression (Drisdelle & Eimer, 2021; Gaspelin & Luck, 2019; Luck et al., 2021). However, our view is that a suppression process should be classified as “proactive” only if the inhibition process occurs before display onset. Therefore, proactive featural suppression relies on foreknowledge of the to-be-suppressed stimulus’ feature(s), and in other words, the to-be-suppressed stimulus must possess some learnable consistent features. Suppression that can potentially prevent attention to the salient distractor without learnable distractor attributes (Sawaki & Luck, 2010) may also exist, but it requires a different mechanism than proactive suppression, as it is logically impossible for saliency signals to become available before display onset. This point will be further discussed in a later part of the article.

2 As noted by Kerzel & Burra (2020), this P-N flip also seemed to have occurred in Gaspelin & Luck (2018a)’s experiment 3. The ERP waveform is visually inspectable from Figure 9 in Gaspelin & Luck (2018a). However, the second ERP component was not analyzed or discussed in Gaspelin & Luck (2018a).

3 We chose not to simulate the EEG at set size 8 reported by Barras & Kerzel (2016): In their experiments, the target is always a circle among heterogenous distractors that are either a triangle, a square, or a diamond. The circle target may thus be highly salient as the curvilinear singleton. In fact, Gaspelin & Stilwell (2021) reported saliency model estimations on a similar display used in their Experiment 4 and found the circle be even more salient than the color singleton distractor. Having a high-salient search target can greatly impact the search strategy, as it may allow for the use of singleton search mode and may even negate the necessity of engaging in proactive featural suppression. As we wish to focus on EEG from a fixed feature search task, the EEG reported by Barras & Kerzel (2016) is not simulated in this article.

4 Note that a probe array that appears with a 200ms offset probably captures the attentional dynamics at a slightly later timepoint.

